# A novel anaerobic *n*-Hexadecane hydroxylation pathway in *Thermalkanevorax longiformis* gen. nov., sp. nov. isolated from a deep-sea hydrothermal vent

**DOI:** 10.64898/2026.01.09.698600

**Authors:** Huilin Wen, Ge Liu, Chaomin Sun, Rui Liu

**Affiliations:** Laboratory of Experimental Marine Biology & Center of Deep Sea Research, Shandong Province Key Laboratory of Marine Biodiversity and Bio-resource Sustainable Utilization, Institute of Oceanology, Chinese Academy of Sciences, Qingdao, China; Laboratory for Marine Biology and Biotechnology, Qingdao National Laboratory for Marine Science and Technology, Qingdao, China; College of Earth Science, University of Chinese Academy of Sciences, Beijing, China

**Keywords:** *Symbiobacteriaceae*, *n*-Hexadecane, Anaerobic metabolism, Hydroxylation, Hydrothermal environment

## Abstract

Hydrothermal systems contain a wide spectrum of alkanes derived from biological sources and geological processes. However, despite their widespread occurrence in hydrothermal environments, the capacity and mechanisms of anaerobic alkane degradation by members of the phylum *Bacillota* remain poorly understood. In this study, we isolated a bacterial strain (designated strain L01) from the Lost City hydrothermal field (LCHF) by adding crude oil for enrichment culture. Strain L01 can grow anaerobically at 55 °C and pH 7.0 in medium supplemented with *n*-hexadecane, indicating its capability for anaerobic alkane degradation. Phylogenetic analysis based on the 16S rRNA gene sequence (93.11% similarity to its closest described relative) and average amino acid identity (AAI, 66.27-66.58%) revealed that strain L01 represents a novel genus within the family *Symbiobacteriaceae*, for which the name *Thermalkanevorax longiformis* gen. nov., sp. nov. is proposed. Cells of strain L01 are elongated rods measuring approximately 10-20 μm in length and ∼0.2 μm in diameter. Integrated genomic, transcriptomic, and metabolomic analyses revealed a previously unrecognized anaerobic *n*-hexadecane utilization pathway, involving initial alkane activation by AhyA homologous enzymes followed by stepwise oxidation and β-oxidation. Together, these findings provide new insights into anaerobic alkane metabolism by hydrothermal bacteria, expanding current understanding of microbial hydrocarbon degradation in extreme marine environments.

**IMPORTANCE:** Although alkanes have be reported degradation by microorganisms through both aerobic and anaerobic metabolic pathways in hydrothermal ecosystems, the mechanisms underlying anaerobic alkane degradation remain incompletely understood, and the metabolic strategies employed by hydrothermal microorganisms are still poorly characterized. In this study, a novel genus bacterium *Thermalkanevorax longiformis* within the *Symbiobacteriaceae* family was isolated from the LCHF and identified an unrecognized C1-position *n*-hexadecane hydroxylation of AhyA homologous proteins at the anaerobic condition. These findings not only provide fundamental insights into the metabolic strategies of hydrothermal bacteria but also expand our understanding of microbial hydrocarbon degradation in extreme marine environments, offering a potential genomic resource for biotechnological applications in hydrocarbon remediation.

## INTRODUTION

Deep-sea hydrothermal ecosystems are extreme environments formed by the mixing of anoxic, chemically enriched hydrothermal fluids with cold, oxic seawater (1). The serpentinite-hosted Lost City hydrothermal field (LCHF) is a remarkable deep-sea ecosystem, in which geological, chemical, and biological processes are closely interconnected. Seawater circulation through upper mantle peridotites, driven by serpentinization reactions at the LCHF (2), generates highly alkaline hydrothermal fluids (pH 9-11) with temperatures ranging from 40 to 91 °C and enriched in hydrogen, methane, and diverse hydrocarbons (2–6). Hydrothermal ecosystems contain a wide range of alkanes with carbon chain lengths ranging from C9 to C30 (6–8), which can be degraded by microorganisms through both aerobic and anaerobic metabolic pathways (8–10).

Under aerobic conditions, microbial alkane degradation is typically initiated by monooxygenases that catalyze terminal methyl oxidation, sequentially converting alkanes into fatty alcohols, aldehydes, and fatty acids (11, 12). These fatty acids are subsequently funneled into β-oxidation, producing acetyl-CoA that enters the tricarboxylic acid cycle (TCA) for complete mineralization to CO_2_ and H_2_O (12). Multiple enzyme systems have been identified to catalyze the initial hydroxylation of alkanes in bacteria, including propane monooxygenase (C_3_), butane monooxygenase (C_2_-C_9_), CYP153 cytochrome P450 monooxygenases (C_5_-C_12_), AlkB-related non-heme iron monooxygenases (C_3_-C_13_ or C_10_-C_20_), the flavin-binding monooxygenase AlmA (C_20_-C_36_), the flavin-dependent monooxygenase LadA (C_10_-C_30_), and a copper-containing flavin-dependent dioxygenase (C_10_-C_30_) (13).

In the absence of oxygen, alkanes can also be degraded by microorganisms using alternative electron acceptors such as nitrate, sulfate, or ferric iron (14, 15). To date, four oxygen-independent mechanisms for anaerobic alkane activation have been proposed: fumarate addition (16), alkyl-coenzyme M formation (17, 18), hydroxylation (19), carboxylation (20). Among these, fumarate addition is considered the most widespread pathway. For example, *Desulfatibacillum alkenivorans* AK-01, isolated from petroleum-contaminated sendiment, activates alkanes via fumarate addition catalyzed by alkylsuccinate synthase, followed by carbon-skeleton rearrangement, decarboxylation, and β-oxidation, ultimately resulting in complete oxidation to CO_2_ (21). In contrast to fumarate addition, *Candidatus Methanoliparum* isolated from a subsurface oil reservoir performs methanogenic *n*-hexadecane degradation via alkyl-coenzyme M reductase-mediated activation to hexadecyl-CoM, which is further processed through β-oxidation to provide reducing equivalents for methane formation via the Wood-Ljungdahl pathway and canonical methyl-coenzyme M reductase (MCR) (18). In addition, a putative C2-methylene hydroxylase (AhyABC) has been proposed to initiate subterminal hydroxylation of alkanes, forming alkan-2-ols that are subsequently oxidized to ketones, and carboxylated at C3 position. Genes encoding this system and corresponding signature metabolites have been detected in oil reservoir environments (14, 19). Furthermore, strain Hxd3, a sulfate-reducing bacterium isolated from an oil field, degrades alkanes anaerobically via subterminal carboxylation at the C3 position using bicarbonate, coupled with the removal of two adjacent terminal carbon atoms, generating fatty acids one carbon shorter than the alkane. These products may subsequently enter β-oxidation or undergo further transformation through chain elongation and C10 methylation (20, 22).

Despite these advances, the mechanisms underlying anaerobic alkane degradation remain incompletely understood, and the metabolic strategies employed by hydrothermal microorganisms are still poorly characterized. Members of the phylum *Bacillota* are widely distributed in the LCHF (2), but their capacity for anaerobic alkane degradation and the associated molecular mechanisms remain largely unexplored. In this study, we report the isolation of a putative novel genus within the phylum *Bacillota* (class *Clostridia*, order *Eubacteriales*, family *Symbiobacteriaceae*), *Thermalkanevorax longiformis* gen. nov., sp. nov. (strain L01 = MCCC 1K10379), from the Lost City hydrothermal ecosystem. Strain L01 is capable of degrading *n*-hexadecane under anaerobic conditions. However, no similar genes of alkane activation in both aerobic and anaerobic metabolic pathways were detected from the genome of strain L01. By integrating transcriptomic, and metabolomic analyses, this study aims to elucidate the potential molecular mechanisms underlying *n*-hexadecane metabolism in strain L01, thereby providing new insights into anaerobic alkane degradation in hydrothermal and marine microorganisms.

## MATERIALS AND METHODS

### Enrichment, isolation, and culture of strain L01

Enrichment cultures were established in 15 mL Hungate tubes by inoculating 1 g of hydrothermal sediment collected from the Lost City into 10 mL of anaerobic basal medium (BM) supplemented with 1 g of crude oil. The headspace was flushed with a gas mixture of 10% H_2_, 10% CO_2_, and 80% N_2_ to ensure anaerobic conditions. The BM contained (per liter of distilled water): 1.0 g NH_4_Cl, 1.0 g NaHCO_3_, 1.0 g CH_3_COONa, 0.5 g KH₂PO_4_, 0.2 g MgSO_4_·7H_2_O, 25 g NaCl, 0.1 g CaCl_2_, 0.2 g KCl, 1 g yeast extract, 1 g tryptone and 10 mL modified Wolin’s mineral solution (Table S1) (23). The pH was adjusted to 7.0 ± 0.2 prior to autoclaving. Cultures were incubated at 55 °C for 15 days.

After successive transfers, a pure culture was obtained by serial dilution to extinction and designated strain L01. For routine subcultivation, strain L01 was cultivated in basal medium 1 (BM1) supplemented with *n*-hexadecane (>99%) at a final concentration of 0.05% (vol/vol). The BM1 contained (per liter of distilled water): 1.0 g NH_4_Cl, 1.25 g Na_2_CO_3_, 1.0 g CH_3_COONa, 0.5 g KH_2_PO_4_, 0.2 g MgSO_4_·7H_2_O, 25 g NaCl, 0.1 g CaCl_2_, 0.2 g KCl, 1.7 g NaNO_3_, 1 g yeast extract, 2 g tryptone and 10 mL modified Wolin’s mineral solution (Table S1). The pH of the medium was adjusted to 7.0 ± 0.2 prior to autoclaving, and the headspace was flushed with N₂ to maintain anaerobic conditions.

### 16S rRNA gene amplification and phylogenetic analysis

The nearly full-length 16S rRNA gene of strain L01 was amplified using the universal bacterial primers 27F (5’-AGAGTTTGATCCTGGCTCAG-3’) and 1492R (5’-GGTTACCTTGTTACGACTT-3’). PCR amplification was performed in a total reaction volume of 20 μL containing 10 μL of Green Taq Mix (Vazyme, China), 2 μL of genomic DNA template, 1 μL of primer 27F (10-20 pmol), 1 μL of primer 1492R (10-20 pmol), and 6 μL of sterile deionized water. The PCR program consisted of an initial denaturation at 95 °C for 10 min, followed by 30 cycles of denaturation at 95 °C for 15 s, annealing at 54 °C for 15 s, and extension at 72 °C for 25 s. Amplifications were carried out using a Veriti thermal cycler (Applied Biosystems, USA). PCR products were purified and sequenced by Tsingke Biotechnology Co., Ltd (China). The obtained 16S rRNA gene sequence was compared with reference sequences available in the NCBI GenBank database using the BLAST algorithm.

For phylogenetic analysis, the 16S rRNA gene sequence of strain L01 was aligned with those of closely related taxa within the family *Symbiobacteriaceae* retrieved from the NCBI GenBank database using ClustalW implemented in MEGA X software (24). Phylogenetic distances were calculated using the p-distance method, and a neighbor-joining phylogenetic tree was reconstructed. The robustness of the inferred tree topology was evaluated by bootstrap analysis with 1,000 replicates, and bootstrap values greater than 50% were indicated at the corresponding nodes. The 16S rRNA sequences of *Bacillus subtilis* DSM 10 and *Geobacillus stearothermophilus* IFO 12550 were used as outgroup taxa.

To further assess the phylogenetic relationships between strain L01 and related taxa at the whole-genome level, average amino acid identity (AAI) analysis was performed between strain L01 and representatives of the closely related genera *Symbiobacterium, Caldinitratiruptor,* and *Thermaerobacter.* The analyses were conducted using the online tools available on the Majorbio Cloud Platform (China).

### qPCR-based growth curve determination of strain L01

The biomass of strain L01 was quantified by quantitative real-time PCR (qPCR) targeting a strain-specific fragment of the 16S rRNA gene. Each qPCR reaction was performed in a total volume of 10 μL containing 5 μL SYBR^®^ Green Realtime PCR Master Mix (Toyobo, Japan), 3 μL sterile distilled water, 0.5 μL of each primer, and 1 μL of DNA template. Amplification was carried out with an initial denaturation at 95 °C for 1 min, followed by 40 cycles of denaturation at 95 °C for 15 s and annealing/extension at 60 °C for 1 min. A ∼150 bp fragment of the 16S rRNA gene was amplified using primers 16SF (5’-CATCAGGAAATGGGTGTGG-3’) and 16SR (5’-CCTTTCACCCCTGACTTACC-3’). For standard curve construction, the nearly full-length 16S rRNA gene of strain L01 was cloned into the pUCm-T vector (Sangon Biotech, China) and propagated in *Escherichia coli* DH5α. Recombinant plasmids were purified using a Plasmid Extraction Kit (Tiangen, China), and DNA concentrations were determined with a NanoDrop spectrophotometer (IMPLEN, Germany). Plasmid copy numbers were calculated according to the following formula (25):

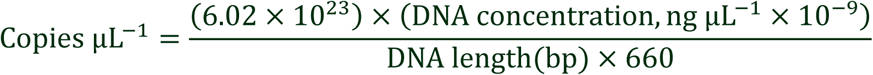

The recombinant plasmid was serially diluted 10-fold to generate eight concentration gradients. Standard curves were constructed by plotting Ct values against the corresponding plasmid copy numbers and were used for absolute quantification. The recombinant plasmid copy number was assumed to be equivalent to the 16S rRNA gene copy number (Supplementary Dataset).

For growth curve determination, 1 mL of bacterial culture was harvested by centrifugation at 7,000 ×*g* for 30 min at 25 °C. The supernatant was discarded, and total genomic DNA was extracted from the cell pellet using a conventional phenol extraction method and resuspended in 50 μL of sterile distilled water (4). The extracted DNA was used as the template for qPCR analysis. And biomass was expressed as 16S rRNA gene copy numbers.

### Genome sequencing and annotation of strain L01

Cells of strain L01 in the exponential growth phase were harvested by centrifugation at 7,000 ×*g* for 30 min at 4 °C. Cell pellets were submitted to Majorbio Co., Ltd. (China) for genomic DNA extraction and whole-genome sequencing. Genome sequencing was performed using a combination of Illumina short-read and PacBio long-read platforms. De novo genome assembly was conducted using Unicycler, incorporating PacBio long reads, followed by polishing with Pilon based on Illumina short-read data. Genome circularization was achieved by trimming terminal overlaps, resulting in a finished chromosomal sequence.

Functional annotation was performed by comparing predicted protein sequences against multiple public databases, including NR, Swiss-Prot, EggNOG, KEGG, Pfam and GO to assign putative gene functions based on sequence similarity.

### Morphological observation

To examine both the overall cellular morphology and ultrastructural features of strain L01, high-resolution images were obtained using transmission electron microscopy (TEM). For whole cell observation, 1 mL of cell suspension was harvested by centrifugation at 7,000 ×*g* for 10 min at 25 °C, washed once with sterile water, and centrifuged again under the same conditions. The resulting pellet was resuspended in approximately 50 μL of sterile water, applied onto copper grids for 30 min, air-dried, and then observed directly using a HITACHI TEM system (HT7700, Hitachi, Japan).

For ultrathin sectioning, 200 mL of cell suspension was collected by centrifugation at 7,000 ×*g* for 15 min at 25 °C. The cell pellet was gently fixed in 1 mL of 2.5% glutaraldehyde (Solarbio, China) at 4 °C overnight, washed three times with 0.1 M phosphate buffer (15 min each), and postfixed with 1% osmium tetroxide for 1-1.5 h. After three additional washes with 0.1 M phosphate buffer (15 min each), cells were dehydrated sequentially in 50%, 70%, and 90% acetone for 15 min each, followed by three changes of 100% acetone for 15-20 min each. Samples were embedded in Epon 812 resin by stepwise infiltration with acetone-resin mixtures (2:1vol/vol for 30 min at 25 °C, 1:2 vol/vol for 1.5 h at 37 °C), followed by pure resin at 37 °C for 2-3 h. Polymerization was carried out sequentially at 37 °C, 45 °C, and 60 °C for 24 h at each temperature. Ultrathin sections (∼70 nm) were prepared using a Reichert-Jung Ultracut E ultramicrotome (Austria), collected on copper grids, and stained with uranyl acetate for 15 min followed by lead citrate for 15 min prior to observation.

For scanning electron microscope (SEM), 20 mL of cell suspension was harvested by centrifugation at 7,000 ×*g* for 10 min at 25 °C and washed with 0.1 M phosphate-buffered saline (PBS). The pellet was fixed in precooled 5% glutaraldehyde at 4 °C for 2 h, followed by three rinses with 0.1 M PBS (10 min each). Samples were dehydrated sequentially in 30%, 50%, 70%, 80%, 90%, and 100% ethanol for 10 min at each step, followed by critical-point drying for approximately 3 h. Dried samples were examined using a field-emission scanning electron microscope (Gemini 500, ZEISS, Germany).

Gram staining was performed using 1 mL of cell suspension harvested by centrifugation at 7,000 ×*g* for 10 min at 25 °C. After removal of most of the supernatant, the concentrated cell suspension was spread onto a clean glass slide, air-dried, and heat-fixed. Smears were stained using commercial Gram staining kit (Qingdao Hope Bio, China) according to the manufacturer’s instructions and examined under a light microscope.

### Fluorescence labeling and confocal microscopic observation of *n*-hexadecane uptake

To examine cellular uptake of *n*-hexadecane, cells of strain L01 were stained with the lipophilic membrane dye Dil (Beyotime, China) and FITC-labeled *n*-hexadecane (Xi’an Qiyue Biology, China), followed by observation using confocal laser scanning microscopy (LSM). Briefly, 1 mL of cell culture was mixed with 1 μL Dil and centrifuged at 7,000 ×*g* for 4 min at 25 °C. The supernatant was discarded, and the cell pellet was resuspended in 1 mL of 0.1 M PBS. Cells were centrifuged again under the same conditions, the supernatant was discarded. And the pellet was resuspended in 1 mL 0.1 M PBS. The resulting cell suspension was mixed with FITC-labeled *n*-hexadecane solution and 1% (wt/vol) agarose at a volume ratio of 10:1:15. The mixture was mounted onto glass slides and examined using a confocal laser scanning microscope (LSM900, ZEISS, Germany).

### Identification of putative anaerobic alkane hydroxylase (AhyA) genes

Putative genes encoding anaerobic alkane hydroxylase (AhyA) were identified from the genome of strain L01 using HMMER (v3.3.1) to search against the CANT-HYD database (26). Hits were filtered using a stringent threshold of full-sequence (E-value < 1×10^−10^) and only sequences meeting this criterion were retained as putative AhyA-encoding genes.

The conserved motifs of AhyA reference sequence from *Desulfococcus oleovorans* Hxd3 (WP_012173623.1) in CANT-HYD database and homologous amino acid sequences from strain L01 were discovered by using a motif-based sequence analysis tool (The MEME Suite, version 5.5.9) (27). The predicted results were further applied to subsequent molecular docking simulations.

### Molecular docking simulation

Based on the screening of anaerobic alkane hydroxylase (AhyA) homologs and transcriptomic analysis in strain L01, three-dimensional (3D) structures of the AhyA protein from *D*. *oleovorans* Hxd3 and eight putative AhyA proteins from strain L01 were obtained in PDB format. Protein structures were retrieved from the SWISS-MODEL server (https://swissmodel.expasy.org/) and the AlphaFold Protein Structure Database (https://alphafold.ebi.ac.uk/). The SMILES structure of ligand (*n*-hexadecane) was obtained from the PubChem database of NCBI (https://pubchem.ncbi.nlm.nih.gov/).

Molecular docking simulations between AhyA proteins and *n*-hexadecane were performed using the SwissDock (28) online service (https://www.swissdock.ch/), which implements the AutoDock Vina docking engine (29). The docking grid box was set at 30 × 30 × 30 to cover two motifs in 3D structures, and the sampling exhaustivity was set to at least 4. The predicted binding affinities (kcal/mol) of different docking conformations were calculated to estimate the binding energy required for protein-ligand complex formation. Lower affinity values indicate more favorable binding conformations and stronger predicted protein-ligand interactions.

### Transcriptome sequencing and differential expression analysis

Strain L01 was cultivated in BM1 medium supplemented with a trace amount of *n*-hexadecane (15 μL per 200 mL medium) for 24 h. Subsequently, the experimental group was further supplemented with an additional 85 μL of *n*-hexadecane, whereas no additional substrate was added to the control group. Cultures from both groups were harvested at 1 h and 4 h after *n*-hexadecane induction. Cells were collected by centrifugation at 7,000 ×*g* for 30 min at 4 °C, and the pellets were submitted to Majorbio Co., Ltd. (Shanghai, China) for total RNA extraction and transcriptome sequencing.

Functional annotation of expressed genes was performed by comparison against multiple databases, including NR, Swiss-Prot, EggNOG, KEGG, Pfam, and GO. Differential gene expression analysis between experimental and control groups was conducted using DESeq2. All transcriptomic experiments were performed with three biological replicates.

### Determination of *n*-hexadecane degradation

For the degradation assay, 5 μL of *n*-hexadecane was added to 10 mL of BM1 medium. Abiotic controls were prepared without inoculation, while experimental cultures were inoculated with strain L01 at 4% (vol/vol). After incubation, residual *n*-hexadecane was extracted by adding 2 mL of *n*-hexane to each culture, followed by overnight extraction. The upper organic phase was collected for analysis. Quantification of *n*-hexadecane was performed using an Agilent 8860 gas chromatograph equipped with a flame ionization detector (FID) and an HP-5 capillary column (30 m × 320 μm × 0.25 μm; Agilent Technologies, USA). Analytical-grade *n*-hexadecane (≥99.5%, Macklin, China) dissolved in chromatographic-grade *n*-hexane (≥99%, Macklin, China) was used to prepare standard solutions. Nitrogen was used as the carrier gas. Samples were injected in split mode (split ratio 10:1) at an inlet temperature of 200 °C with a split flow rate of 65 mL/min. The oven temperature program was as follows: 70 °C for 0.5 min, ramped to 190 °C at 20 °C/min, and held for 6.25 min. The FID temperature was set at 250 °C, and the injection volume was 1 μL. A calibration curve was generated by analyzing serially diluted *n*-hexadecane standards (0-11997.77 μmol/L) under identical chromatographic conditions, and linear regression was performed using peak area versus concentration. Quantification of extracted samples was conducted using the external standard method. All measurements were performed in triplicate.

The *n*-hexadecane degradation rate was calculated using the following equation (30):

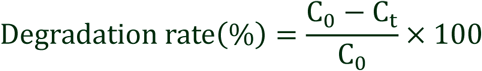

where C_0_represents the mean *n*-hexadecane concentration in abiotic controls averaged across all sampling time points, and C_t_ represents the residual *n*-hexadecane concentration in inoculated cultures at time *t*. This normalization approach was applied to minimize variability caused by volatilization, sampling loss, and extraction-related fluctuations.

### Metabolomic analysis following *n*-hexadecane addition

Both control and experimental groups were cultured in BM1 medium supplemented with 1 g/L succinate for 2 days. Subsequently, 100 μL of *n*-hexadecane was added to 200 mL of culture in the experimental group, whereas no substrate was added to the control group. After an additional 6 h of incubation, cells were harvested by centrifugation at 7,000 ×*g* for 30 min at 4 °C. The supernatant was discarded, and cell pellets were washed twice with 0.1 M PBS and centrifuged at 7,000 ×*g* for 5 min at 4 °C. The resulting pellets were submitted to Novogene Co., Ltd. (Beijing, China) for untargeted metabolomic analysis.

Metabolites were extracted by resuspending the cell pellets in 300 μL of 80% (vol/vol) methanol. Samples were flash-frozen in liquid nitrogen for 5 min, thawed on ice, vortexed for 30 s, and ultrasonicated for 6 min. After centrifugation at 3,000 ×*g* for 1 min at 4 °C, the supernatants were collected and lyophilized. The dried extracts were reconstituted in 70 μL 10% (vol/vol) methanol and subjected to LC-MS analysis (31, 32).

UHPLC-MS/MS analyses were performed using a Vanquish UHPLC system (Thermo Fisher Scientific, Germany) coupled to an Orbitrap Q Exactive™ HF-X mass spectrometer (Thermo Fisher Scientific, Germany). Chromatographic separation was achieved on a Hypersil Gold column (100 × 2.1 mm, 1.9 μm) at a flow rate of 0.2 mL/min using a 12-min linear gradient. The mobile phases consisted of eluent A (0.1% formic acid in water) and eluent B (methanol) for both positive and negative ion modes. The gradient program was as follows: 2% B for 1.5 min, 2% to 85% B for 3 min, 85% to 100% B for 10 min, 100% to 2% B at 10.1 min, and 2% B at 12 min. The mass spectrometer was operated in both positive and negative ion modes with a spray voltage of 3.5 kV, capillary temperature of 320 °C, sheath gas flow rate of 35 psi, auxiliary gas flow rate of 10 L/min, S-lens RF level of 60, and auxiliary gas heater temperature of 350 °C.

Raw UHPLC-MS/MS data were processed using XCMS for peak detection, retention time alignment, and quantification. Metabolites were identified by comparison with a self-built high-quality MS/MS spectral database (NovoMetDB), based on adduct ions and a mass tolerance of 10 ppm. Background ions were removed using blank samples. Quantitative data were normalized as follows: Relative peak areas = Raw quantitative value of samples/ (The sum of quantitative value of samples/ The sum of quantitative value of QC). Metabolites with coefficients of variation (CVs) greater than 30% in quality control (QC) samples were excluded. The remaining metabolites were retained for downstream analysis and annotated using the KEGG, HMDB, and LIPID MAPS databases.

### Data availability

The 16S rRNA and genome sequences of strain L01 have been deposited in the NCBI database under accession number PX804844 and PRJNA1347602. The raw transcriptomic sequencing data have been deposited in the NCBI Sequence Read Archive under the same BioProject accession number (PRJNA1347602).

## RESULTS AND DISCUSSION

### Hydrocarbons-mediated isolation and cultivation of hydrothermal microorganisms

In the Lost City hydrothermal field and other regions along the Mid-Atlantic Ridge, hydrocarbons of varying chain lengths have been reported with a relatively high abundance (33), including C_1_-C_4_ short-chain hydrocarbons (34), C_9_-C_14_ aliphatic hydrocarbons (6), C_6_-C_16_ aromatic hydrocarbons in fluids (35), and long-chain alkanes C_15_-C_30_ in serpentines (5). Despite their abundance, only a few anaerobic alkane-degrading microorganisms have been successfully isolated and cultured from these environments. To enrich microorganisms capable of anaerobic alkane utilization from the Lost City hydrothermal field, we collected hydrothermal sediment samples using the Jiaolong Human-Operated Vehicle (China) and incubated them anaerobically in 10 mL of BM medium supplemented with 1 g of crude oil at 55 °C. As crude oil containing a complex mixture of hydrocarbons (36, 37), it was used in the enrichment process to promote the growth of alkane-degradation microorganisms. After 15 days of enrichment, a bacterial strain was found to dominate the culture. This strain, designated L01, was subsequently isolated in pure culture through successive passage and medium optimization. These results indicate that the addition of crude oil or other hydrocarbons can facilitate the isolation and cultivation of alkane-degrading microorganisms from hydrothermal environments, potentially including strains that have never been cultured before (4).

### Strain L01 as a representative of potential novel genus in the family *Symbiobacteriaceae*

Analysis of the 16S rRNA sequence (1,444 bp) of strain L01 revealed that it belongs to the family *Symbiobacteriaceae* and shares 93.11% sequence identity (the highest in the NR database of NCBI) with *Symbiobacterium ostreiconchae* JCM 15048 (AP042454.1). Phylogenetic analysis was conducted based on 16S rRNA sequences of strain L01 and other species in the family *Symbiobacteriaceae*. Because the family currently comprises few recognized species, several representatives from other families were also included. Although strain L01 clustered as a sister clade with members of the genus *Symbiobacterium*, it formed an independent branch within this lineage and was clearly separated from the genus *Caldinitratiruptor* (Fig. 1A). These observations suggest that strain L01 may represent a novel species within a potential novel genus. To further evaluate its genomic relatedness, average amino acid identity (AAI) values were calculated between strain L01 (Fig. S2) and representative genomes in *Symbiobacteriaceae* family. Strain L01 shared only 66.27-66.58% identities with recognized *Symbiobacterium* species (Fig. 1B), well below the threshold typically associated with species-level delineation and near the lower boundary proposed for genus-level affiliation (38). Moreover, its AAI values with neighboring genera (Fig. 1B), including *Caldinitratiruptor* and *Thermaerobacter*, were even lower (approximately 53.56-55.37%), whereas genus boundaries are generally defined within the 60-80% AAI range (38). Collectively, the deep branching observed in the 16S rRNA phylogeny and the low intergenomic AAI values indicate that strain L01 represents a distinct and deeply divergent lineage, consistent with a previously unrecognized genus-level taxonomic position. Therefore, these results strongly support classifying strain L01 as the type strain of a novel species in a novel genus, for which the name *Thermalkanevorax longiformis* gen. nov., sp. nov. is proposed. *Thermalkanevorax longiformis* gen. nov., sp. nov. (type strain L01 = MCCC 1K10379) has been deposited in the Marine Culture Collection of China (MCCC).

**Fig 1.**
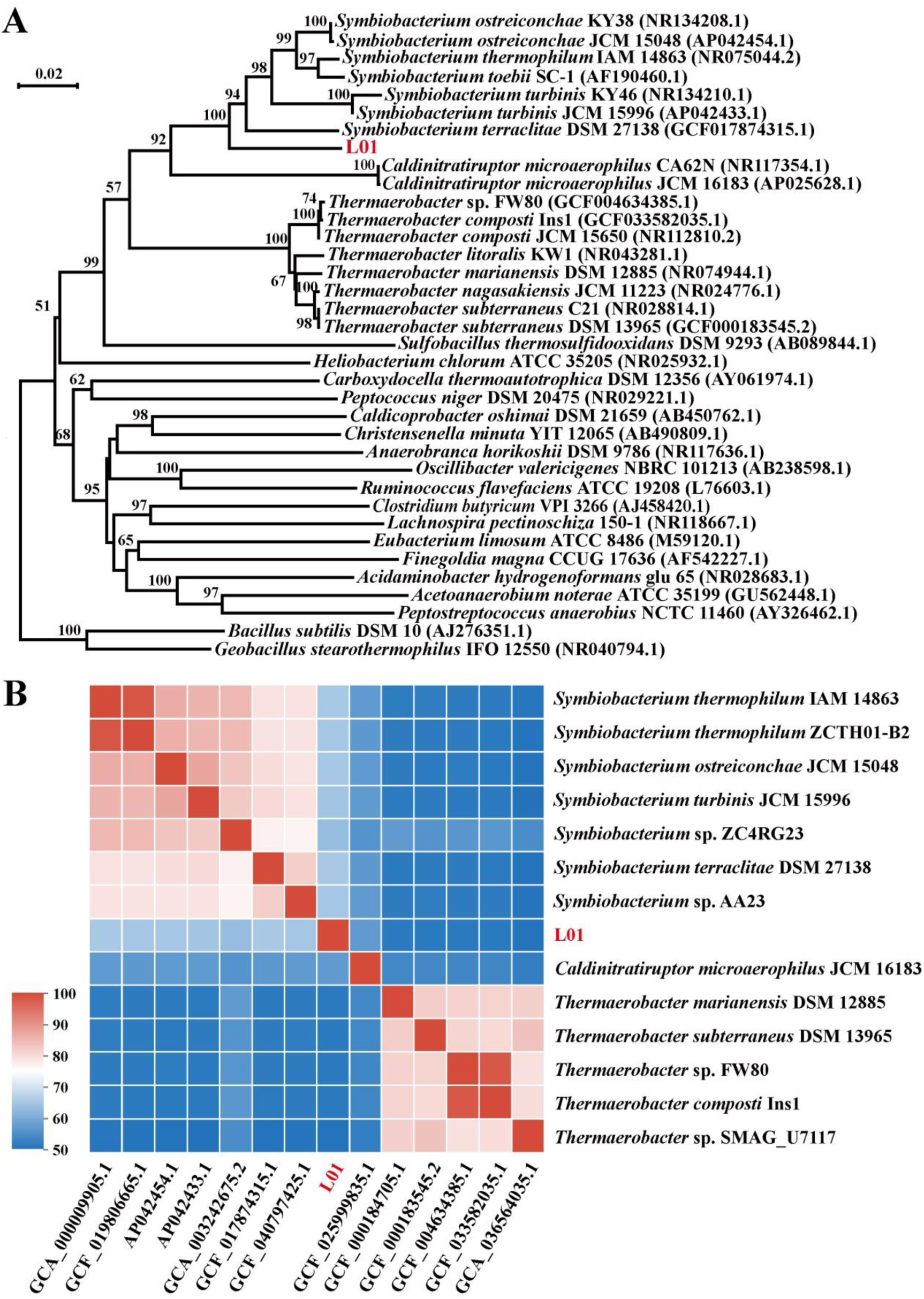
Phylogenetic relationships of strain L01. (A) Neighbor-joining phylogenetic tree based on 16S rRNA sequences showing the phylogenetic position of strain L01 among members of the family *Symbiobacteriaceae*. Bootstrap values (≥50%) based on 1,000 replicates are indicated at branch nodes. *Bacillus subtilis* DSM 10 and *Geobacillus stearothermophilus* IFO 12550 were used as outgroup taxa. (B) The heatmap showing average amino acid identity (AAI) values among strain L01 and representative genomes of the genus *Symbiobacterium*, *Caldinitratiruptor* and *Thermaerobacter*. Colors indicate the degree of amino acid identity, with red representing higher similarity. The clear separation of L01 from described *Symbiobacterium* species and closely related genera at the AAI level supports its placement as a deeply divergent lineage at the genus level.

Morphological characterization via TEM revealed that cells of strain L01 are slender, elongated rods, approximately 10-20 μm in length and ∼0.2 μm in diameter (Fig. 2B, D). Occasional observation of a single terminal flagellum suggests that L01 is motile, a trait further supported by SEM, which revealed numerous flagellum-like appendages distributed around both individual cells and aggregates (Fig. 2G-I). This motility likely facilitates chemotactic movement toward hydrocarbons and direct contact with substrates in the environment. Notably, both TEM and SEM analyses consistently showed frequent associations with neighboring cells, indicating a strong tendency for aggregation and potential cooperative behavior (Fig. 2A, C, G). Ultrathin-section TEM further revealed electron-dense intracellular structures and intimate connections between adjacent cells, suggesting complex cellular interactions (Fig. 2E, F). Interestingly, upon entering the stationary growth phase, strain L01 occasionally formed necklace-like filamentous chains composed of serially aligned spherical bodies (Fig. S1). This feature resembles the endospore-like cellular structure reported in *S*. *thermophilum*, a member of the same family (39). Given the ability of strain L01 to grow at elevated temperatures and the occurrence of similar thermophilic spores in deep-sea hydrocarbon-associated environments (40), these structures may represent an ecological adaptation to thermal or hydrocarbon stress (Table S2). However, further physiological and genetic evidence still is required to confirm their biological function.

**Fig 2.**
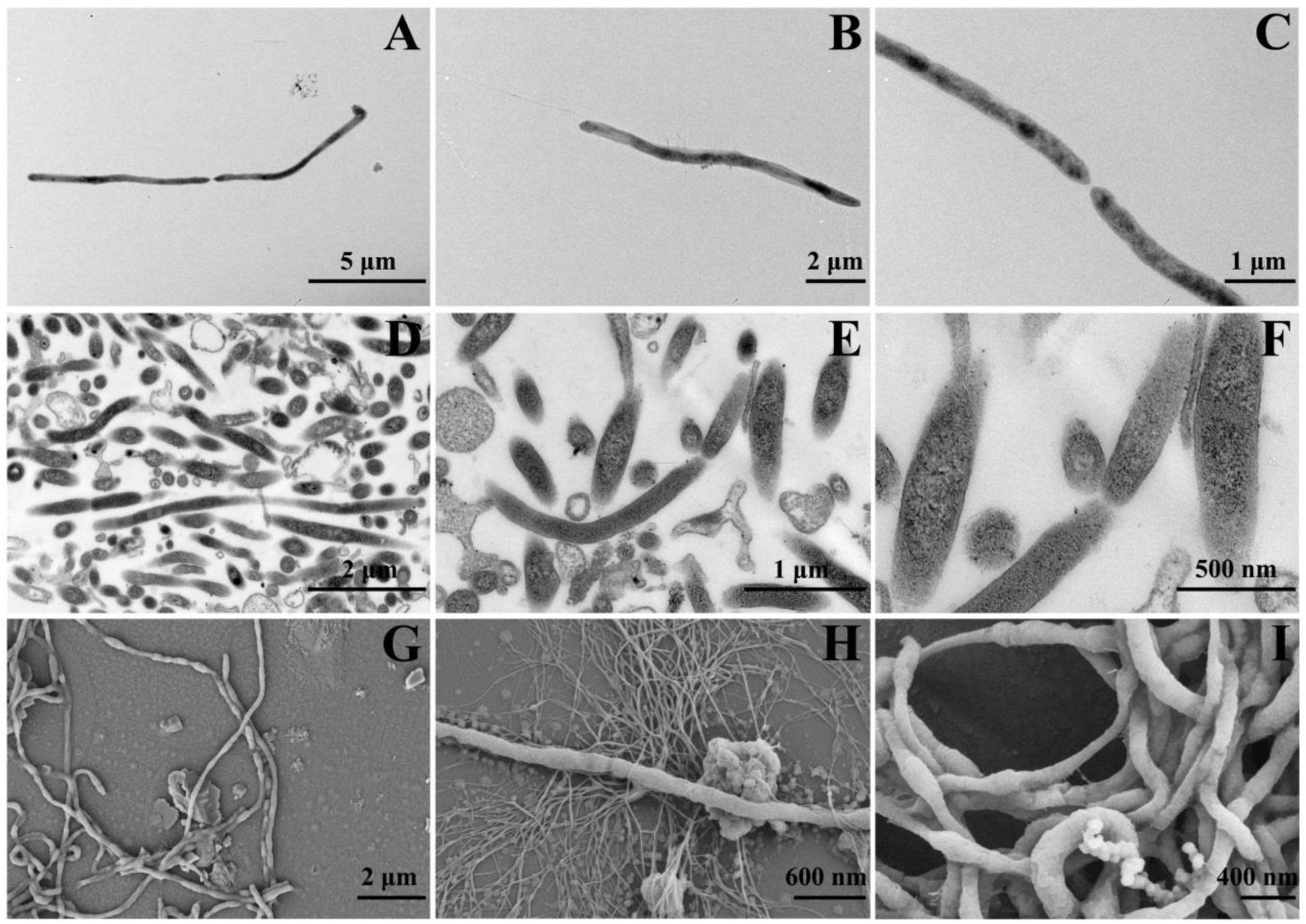
Microscopic observations of strain L01 using transmission electron microscopy (TEM) and scanning electron microscopy (SEM). (A-C) TEM images showed that cells of strain L01 are slender rods with a single terminal flagellum. And cells frequent close contact or potential physical connections between neighboring cells. (D-F) Ultrathin-section TEM images revealed the intracellular structures of strain L01. (G-I) SEM images showed that cells form intertwined filamentous networks with extensive physical contacts. Numerous filamentous or flagellum-like structures were distributed on the cell surface and within cell aggregates (H).

### *n*-Hexadecane promotes the growth of strain L01

Due to the high viscosity of crude oil, direct assessment of the growth of strain L01 was severely impeded. Therefore, alkanes of different chain lengths were evaluated as substitutes, and *n*-hexadecane was identified as the only compound that exerted a growth-promoting effect comparable to that of crude oil. This observation suggests that *n*-hexadecane is likely a key component of crude oil responsible for stimulating the growth of strain L01. To quantitatively assess the effect of *n*-hexadecane on the growth of strain L01, the 16S rRNA gene copy number was used as a proxy for biomass accumulation (Fig. S3). In the presence of *n*-hexadecane, strain L01 entered the exponential growth phase between 6 and 18 h and reached the stationary phase at approximately 24 h; with 16S rRNA gene copy numbers peaking between 18 and 24 h, at ∼4×10^9^ copies/mL (Fig. 3A). In contrast, in the absence of *n*-hexadecane, both the onset of exponential growth and entry into the stationary phase were markedly delayed, and the maximum gene copy number reached only ∼1.6×10^9^ copies/mL at 48 h. These growth profiles clearly demonstrate that *n*-hexadecane significantly enhances the growth of strain L01. The rapid growth observed in the presence of *n*-hexadecane further suggests strain L01 can efficiently access, transport, and metabolize this carbon source for energy generation. Next, the *n*-hexadecane degradation rate by strain L01 was quantified by gas chromatography using an external calibration curve (Fig. S4). Over a 10-day incubation period, the concentration of *n*-hexadecane decreased continuously, with the degradation rate increasing from 7.3% on day 1 to 24.2% on day 10 (Fig. 3B). In parallel, 16S rRNA gene copy numbers were measured at each time point to estimate biomass accumulation.

**Fig 3.**
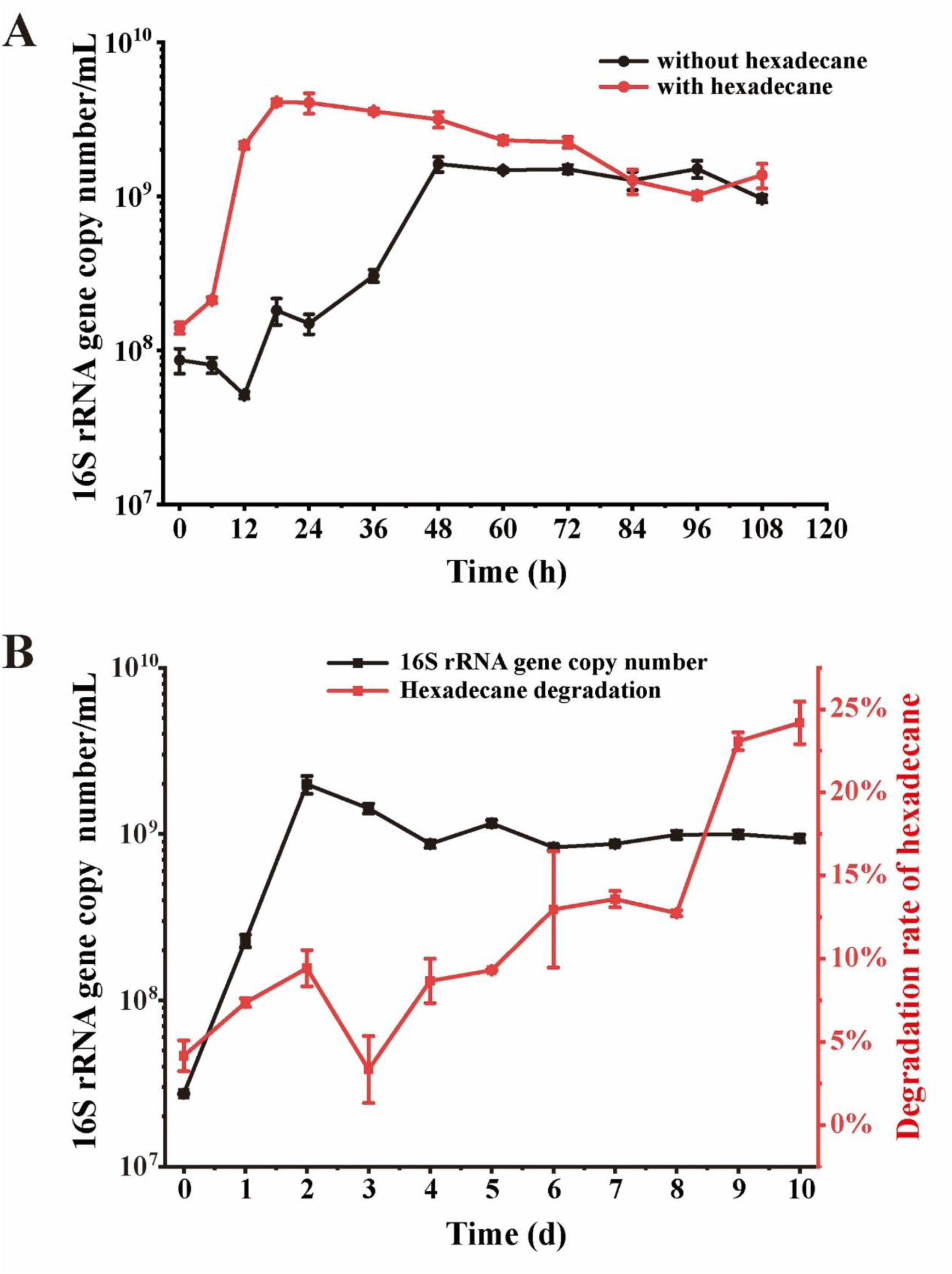
Growth characteristics of strain L01 and *n*-hexadecane degradation. (A) Growth curve of strain L01 under *n*-hexadecane-supplemented and unsupplemented conditions. (B) Growth curve of strain L01 cultivated in hexadecane-supplemented medium and hexadecane degradation rate by strain L01 during cultivation.

Compared with the growth dynamics shown in Fig. 3A, the exponential and stationary phases were shifted to later time points in this experiment. This difference is most likely attribute to variations in initial inoculum density and culture volume between the two experimental setups. Specifically, the larger culture volume (200 mL) and higher initial cell abundance likely facilitated faster population expansion and higher biomass accumulation than the small-volume culture (10 mL), consistent with the enhanced growth observed in Fig. 3A (red line).

Uptake of *n*-hexadecane by strain L01 was further examined using confocal laser scanning microscopy. Cell membranes were stained with Dil to visualize cellular morphology, while *n*-hexadecane was labeled with FITC. Dil staining produced a clear red fluorescence outlining the cells (Fig. 4A), whereas the green fluorescence signal from FITC-labeled *n*-hexadecane was closely associated with the cells (Fig. 4B). In the merged image, the clear colocalization of *n*-hexadecane signals with cell membranes was observed (Fig. 4C), indicating that strain L01 is capable of directly taking up this hydrophobic substrate. The elongated cell morphology and potential flagellum-mediated motility may enhance cell–substrate contact, thereby increasing substrate availability. This trait is consistent with the physiological adaptation of strain L01 to utilize long-chain alkanes as carbon source in hydrothermal environments.

**Fig 4.**
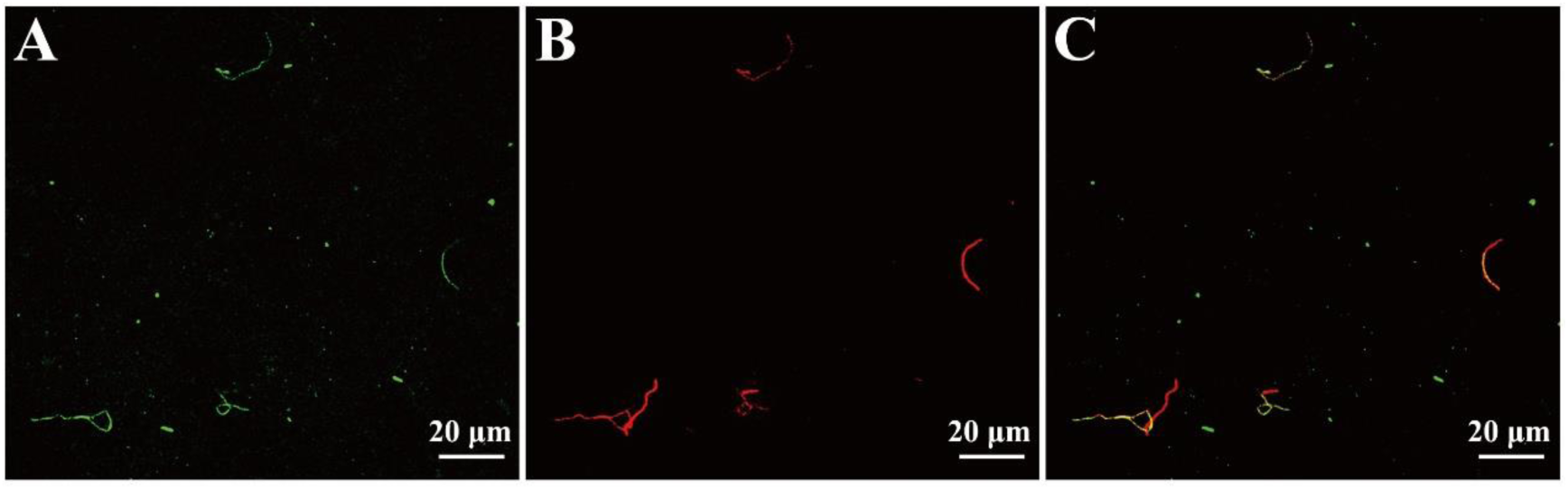
Confocal laser scanning microscopy images showing *n*-hexadecane uptake by strain L01. (A) The signals FITC-labeled *n*-hexadecane in strain L01. (B) Cells were stained with Dil (red) to visualize the cell membrane. (C) The merged image shows colocalization of *n*-hexadecane with the cellular signal. Scale bars, 20 μm.

### A novel *n*-hexadecane metabolic pathway in strain L01

Strain L01 harbors a single circular chromosome of 4.3 Mbp with a GC content of 65.75%, and no plasmid was detected. The circular genome map is shown in Fig. S2. Compared with *Symbiobacterium thermophilum* reported in 2004 (3.57 Mbp; GC content, 68.7%), strain L01 possesses a larger genome but a slightly lower GC content. Genome annotation identified a total of 41 genes associated with sporulation (Table S2), which is consistent with the endospore-like structures observed by TEM. Genome annotation revealed that strain L01 lacks key genes previously reported to be involved in the initial activation of long-chain alkanes under either aerobic or anaerobic conditions, including *alkB*, *almA*, *ladA*, and *assA* (13, 21). This absence suggests that L01 may employ alternative enzymatic strategy for alkane activation. To further explore its alkane degradation potential, HMMER (version 3.3.1) searches were performed against the CANT-HYD database (26). Interestingly, most of genes associated with alkane hydroxylation were not identified in strain L01, excepting 8 genes encoding putative anaerobic alkane C2-methylene hydroxylase (AhyA) (Table S3). These homologous proteins were found to share two conserved motifs with the reference AhyA protein from *D*. *oleovorans* Hxd3 (*Do*AhyA). Molecular docking simulations further demonstrated that these two conserved motifs cooperatively participate in the binding of *n*-hexadecane (Fig. S5 and S6). The results might suggest that AhyA homologous proteins were key molecules responsible for *n*-hexadecane hydroxylation in strain L01. In addition, the genome of strain L01 includes a complete fatty acid β-oxidation and tricarboxylic acid (TCA) pathways, enabling the metabolism of alkane-derived intermediates (Supplementary Dataset). Collectively, these genomic features indicate that strain L01 possesses the genetic potential for anaerobic alkane degradation via an alternative activation mechanism.

To further elucidate the metabolic pathway of *n*-hexadecane degradation in strain L01, transcriptomic analyses were conducted to refine candidate genes identified from genome-based predictions. After 1 h of *n*-hexadecane induction, 26 genes were significantly up-regulated and 2 genes were down-regulated (Fig. S7). In contrast, after 4 h of induction, the number of differentially expressed genes increased markedly, with 534 genes up-regulated and 57 genes down-regulated (Fig. S8), indicating a progressively intensified transcriptional response to *n*-hexadecane exposure. Several AhyA homologous coding genes potentially involved in the initial hydroxylation of *n*-hexadecane were up-regulated at both 1 h and 4 h (Fig. 5A). Notably, genes 0944 and 4094 were significantly induced at 1 h, with gene 0944 exhibiting the highest fold change (log_2_FC=1.8). At 4 h, genes 3275, 0944, 1087, 1934, and 3382 showed pronounced up-regulation. Previous studies have demonstrated that alkane hydroxylases catalyze the initial hydroxylation step during anaerobic alkane degradation (19). Therefore, these AhyA homologous coding genes are possible to play key roles in the anaerobic activation of *n*-hexadecane in strain L01. In addition, gene 2027 (encoding an alcohol dehydrogenase) and gene 3698 (encoding an aldehyde dehydrogenase) was significantly up-regulated at 1 h and 4 h, respectively (Fig. 5A). These expression patterns are consistent with a stepwise oxidation process in which 1-hexadecanol is converted to hexadecanal and subsequently oxidized to hexadecanoic acid. To investigate the downstream metabolism of hexadecanoic acid, transcriptional changes in genes associated with fatty acid β-oxidation and the TCA cycle were examined. Coding genes involved in β-oxidation, including *fadD* (long-chain fatty acid-CoA ligase), *acd* (acyl-CoA dehydrogenase), *crt* (enoyl-CoA hydratase), *lcdH* (3-hydroxyacyl-CoA dehydrogenase), and *fadA* (acetyl-CoA acyltransferase), were all up-regulation at both 1 h and 4 h, with generally higher fold changes at 4 h (Fig. 5B). This result indicates that hexadecanoic acid may be primarily product upon prolonged *n*-hexadecane activation and funneled into β-oxidation to generate acetyl-CoA. Coding genes involved in the TCA cycle were also up-regulated (Fig. 5B). For instance, *gltA* (citrate synthase) and *korB/korA* (2-oxoglutarate ferredoxin oxidoreductase) showed pronounced induction at both time points, with higher expression levels at 4 h. Meanwhile, *sdhB* (succinate dehydrogenase), *fumB* (fumarate hydratase), and *ldh* (malate dehydrogenase) showed moderate up-regulation at both 1 h and 4 h, whereas *IDH3* exhibited only modest induction. Overall, coding genes involved in both β-oxidation and the TCA cycle exhibited stronger induction at 4 h than at 1 h, indicating a sequential regulatory cascade in response to *n*-hexadecane metabolism. *n*-Hexadecane activation likely increases intracellular levels of hexadecanoic acid, thereby stimulating β-oxidation, β-oxidation-derived acetyl-CoA subsequently enhances flux through the TCA cycle.

**Fig 5.**
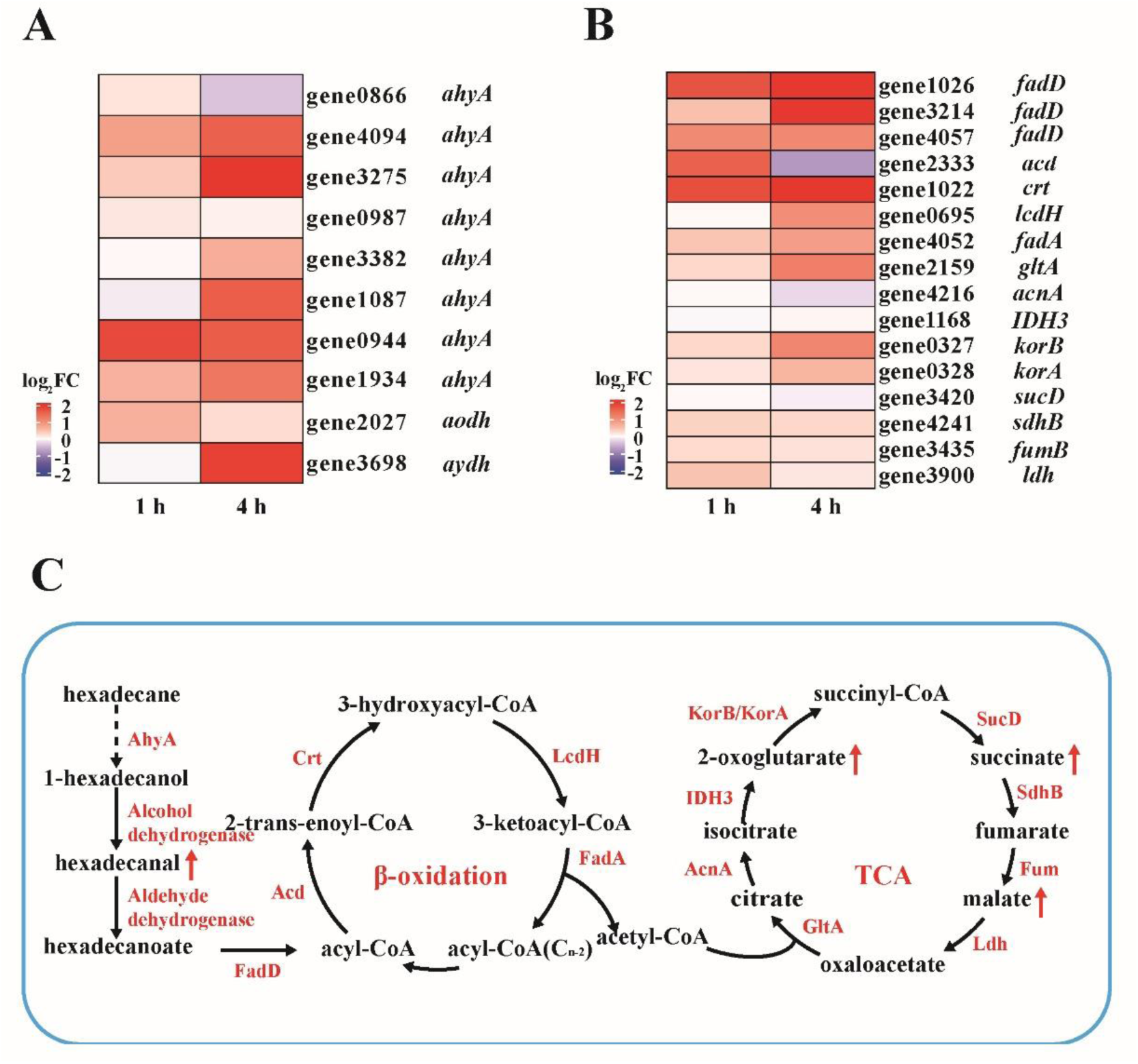
Coding gene expression profiles and proposed metabolic pathway for *n*-hexadecane degradation in strain L01. (A) A heatmap of differentially expressed coding genes involved in the initial oxidation of *n*-hexadecane, including *ahyA* (putative alkane hydroxylase), *aodh* (alcohol dehydrogenase), and *aydh* (aldehyde dehydrogenase). (B) A heatmap of differentially expressed coding genes involved in β-oxidation, including *fadD* (long-chain fatty acid-CoA ligase), *acd* (acyl-CoA dehydrogenase family protein), *crt* (enoyl-CoA hydratase), *lcdH* (3-hydroxyacyl-CoA dehydrogenase), and *fadA* (acetyl-CoA acyltransferase), as well as coding genes involved in the tricarboxylic acid (TCA) cycle, including *gltA* (citrate synthase), *acnA* (aconitate hydratase), *IDH3* (isocitrate dehydrogenase), *korA/korB* (2-oxoglutarate ferredoxin oxidoreductase), *sucD* (succinyl-CoA synthetase), *sdhB* (succinate dehydrogenase), *fumB* (fumarate hydratase), and *ldh* (malate dehydrogenase). Gene expression was measured at 1 h and 4 h following *n*-hexadecane induction. (C) Proposed metabolic pathway for *n*-hexadecane degradation. *n*-Hexadecane is initially oxidized to hexadecanoate, which is subsequently degraded via β-oxidation to produce acetyl-CoA that enters the TCA cycle. Genes encoding the enzymes shown in red are upregulated during *n*-hexadecane induction. Metabolites indicated by upward arrows were upregulated in metabolomic profiles following *n*-hexadecane supplementation.

To validate the proposed metabolic pathway in strain L01, comparative metabolomic analyses were performed using six biological replicates, in which cells were initially cultured in succinate-supplemented basal medium for 48 h and subsequently induced with *n*-hexadecane for 6 h, with non-induced cultures serving as controls. Hexadecanal showed an overall increase in the hexadecane-induced group, supporting its role as a key intermediate in *n*-hexadecane degradation. Based on the metabolite profiles, *n*-hexadecane is likely first oxidized to 1-hexadecanol, then converted to hexadecanal, and subsequently oxidized to hexadecanoic acid, which enters the β-oxidation pathway. Although hexadecanoic acid exhibited a decreasing trend in the induced group, increased levels of tetradecanoic acid and dodecanoic acid were detected, indicating progressive chain shortening via successive β-oxidation cycles. In addition, elevated levels of 12-hydroxydodecanoic acid were observed, accompanied by increases in dodecanedioic acid, sebacic acid, suberic acid, and adipic acid. These results suggest that dodecanoic acid may undergo cytochrome P450-mediated hydroxylation to form 12-hydroxydodecanoic acid, which is subsequently converted to dodecanedioic acid and further processed through multiple rounds of β-oxidation, yielding shorter-chain dicarboxylic acids. Furthermore, several intermediates in TCA cycle were detected changes in the abundance by metabolomic analysis (Supplementary Dataset). Specifically, 2-oxoglutarate and succinate were significantly increased, malate showed an overall increasing trend, whereas fumarate and aconitic acid showed an overall decreasing trend. The reduction in aconitic acid may reflect enhanced metabolic flux toward 2-oxoglutarate, while the concomitant increase in malate and decrease in fumarate are likely associated with accelerated fumarate consumption.

Based on integrated genome, transcriptome, and metabolome analyses, strain L01 is proposed to degrade *n*-hexadecane under anaerobic conditions via the pathway illustrated in Fig. 5C. *n*-Hexadecane is initially activated through hydroxylation at the C1 position by an AhyA-like enzyme, generating a hydroxylated intermediate. This intermediate is subsequently converted to hexadecanal by an alcohol dehydrogenase, and further oxidized to hexadecanoic acid by an aldehyde dehydrogenase. Hexadecanoic acid is then metabolized through β-oxidation, generating acetyl-CoA that enters the TCA cycle to support energy generation and biomass formation. Previous studies have reported that anaerobic alkane hydroxylation typically occurs at the C2 position, catalyzed by enzymes such as AhyA and EBDH-like enzymes (14, 27). Notably, our results suggest that *n*-hexadecane activation in strain L01 may involve hydroxylation at the C1 position even under anaerobic conditions. This reaction is most likely catalyzed by a previously uncharacterized AhyA-homologous enzyme, indicating a potentially alternative mode of alkane activation in anaerobic microorganisms. Nevertheless, direct molecular and biochemical evidence confirming the enzymatic activities proposed herein remains limited. Future investigations focusing on gene functional validation and systematic enzymatic characterization are essential to substantiate the hypothesized metabolic mechanism.

## CONCLUSIONS

In this study, we isolated a novel anaerobic thermophilic bacterium, *Thermalkanevorax longiformis* gen. nov., sp. nov. (strain L01), from the Lost City hydrothermal field. Phylogenetic and genomic analyses confirmed its status as a distinct genus within the *Symbiobacteriaceae* family, exhibiting unique morphological features as slender, elongated rods. Combining genomic, transcriptomic, and metabolomic data, we discovered a previously unrecognized anaerobic *n*-hexadecane utilization pathway, likely initiated by AhyA-homologous enzymes followed by downstream β-oxidation and TCA cycle. These findings not only provide fundamental insights into the metabolic strategies of hydrothermal bacteria but also expand our understanding of microbial hydrocarbon degradation in extreme marine environments, offering a potential genomic resource for biotechnological applications in plastic and hydrocarbon remediation.

### Description of *Thermalkanevorax* gen. nov

*Thermalkanevorax* (Ther.mal.ka.ne.vo’rax. Gr. fem. n. thermē, heat; N.L. neut. n. alkanum, alkane; L. masc. adj. vorax, devouring; N.L. masc. n. *Thermalkanevorax*, a heat-loving, alkane-devouring bacterium, referring to its thermophilic lifestyle and ability to utilize linear alkanes).

Cells are Gram-stain-positive (Fig. S9), rod-shaped, and grow under anaerobic conditions. Members of the genus are moderately thermophilic, members of the genus can grow at neutral pH. Linear alkanes can be utilized as carbon and energy sources.

Phylogenetically, the genus *Thermalkanevorax* forms a distinct and deeply branching lineage within the family *Symbiobacteriaceae*, clearly separated from all currently described genera based on 16S rRNA gene phylogeny and average amino acid identity (AAI) analyses.

The genomic DNA G+C content of the type species is 65.75 mol%.

The type species is *Thermalkanevorax longiformis* sp. nov.

### Description of *Thermalkanevorax longiformis sp. nov*

*Thermalkanevorax longiformis* (lon.gi.for’mis. L. adj. *longiformis*, long-shaped, referring to the elongated cell morphology).

Shows the following characteristics in addition to those given for the genus. Cells are Gram-stain-positive rods (Fig. S9), measuring approximately 10-20 μm in length and ∼0.2 μm in diameter. Growth occurs under anaerobic conditions, at 55 °C and at pH 7.0. *n*-Hexadecane can be utilized as a carbon and energy source.

The genomic DNA G+C content of the type strain is 65.75 mol%.

The type strain, L01 (MCCC 1K10379) was isolated from hydrothermal samples collected from the Lost City hydrothermal field.

## ACKNOWLEDGEMENTS

This work was supported by the NSFC Grant (Nos. 42221005, 42530409), Science and Technology Innovation Project of Laoshan Laboratory (Grant Nos. LSKJ202203103 and 2022QNLM030004-3), Shandong Provincial Natural Science Foundation (Grant No. ZR2024ZD49), Major Research Plan of the National Natural Science Foundation (Grant No. 92351301), Qingdao West Coast New District University Presidents Fund (Grant No. E42424101N), Taishan Scholars Program (Grant No. tstp20230637).

## AUTHOR CONTRIBUTIONS

H.W., R.L., and C.S. conceived the study and wrote the manuscript. H.W. and R.L. carried out most of cultivation and culture-based experiments. R.L. and H.W. conducted the bioinformatics analyses. G.L. carried out the venting fluids sampling. All authors have read and approved the manuscript submission.

## ETHICS STATEMENT

This study has no animal or human experiments. There are no ethical issues involved.

